# Species and sex divergence in vocalizations between hybridizing role-reversed shorebirds, Northern Jacana *(Jacana spinosa)* and Wattled Jacana *(Jacana jacana)*

**DOI:** 10.1101/757336

**Authors:** Evan J. Buck, Toni Brown, Gina Zwicky, Elizabeth P. Derryberry, Sara E. Lipshutz

## Abstract

Species-specific vocalizations can act as a reproductive isolating mechanism between closely related populations. We analyzed vocal divergence between two hybridizing species of sex-role reversed polyandrous shorebirds, the Northern Jacana (*Jacana spinosa*) and Wattled Jacana (*Jacana jacana*). We found that *J. spinosa* calls have higher peak frequency and fundamental frequency than *J. jacana* calls. We also compared calls between males and females, as both jacana species are sex-role reversed and females compete for male mates. Males produce calls with a higher peak frequency, exhibit shorter note lengths and emit a greater number of notes within a calling bout than females, which could relate to mate attraction. These results suggest that vocal divergence could act as a behavioral barrier to limit hybridization between the species and vocalizations may function differently between male and female jacanas.

## Resumen. Divergencia específica y sexual en las vocalizaciones de las aves costeras de roles sexuales invertidos Jacana Norteña (*Jacana spinosa*) y Jacana Carunculada (*Jacana jacana*)

Las vocalizaciones especie-específicas pueden actuar como mecanismos de aislamiento reproductivo entre poblaciones de especies estrechamente relacionadas. Analizamos la divergencia en vocalizaciones entre dos especies de aves costeras poliándricas de rol sexual invertido, Jacana Norteña (*Jacana spinosa*) y Jacana Carunculada (*Jacana jacana*). Encontramos que los llamados de *J*. *spinosa* contienen frecuencias pico y fundamental más altas que los llamados de *J*. *jacana*. También comparamos los llamados entre machos y hembras en ambas especies, ya que ambas tiene el rol sexual invertido y las hembras compiten por parejas. Los machos producen llamados con una frecuencia pico mayor, exhiben longitudes menores de notas y emiten un número mayor de notas dentro de un despliegue de vocalizaciones y producen notas de menor duración que las hembras, lo que podría relacionarse con atracción de pareja. Estos resultados sugieren que la divergencia en vocalizaciones podría actuar como barrera comportamental para limitar la hibridación entre las especies y estas vocalizaciones pueden funcionar distintamente entre machos y hembras de jacanas. Estudios futuros utilizando experimentos de reproducción de audio podrían poner a prueba estas hipótesis.

Palabras clave: llamada, hibridación, jacanas, diferencias de sexo, ave costera, divergencia vocal Between closely related species, divergence in mating signals can facilitate reproductive isolation and drive the process of speciation (Irwin and Price 1999; Coyne and Orr 2004). Whereas mating signals are used to attract and compete for mates within populations, divergence between populations can lead to a breakdown in communication such that individuals do not recognize potential mates or rivals (Coyne and Orr 2004). Evidence from a wide range of taxa suggests that the degree of divergence in mating signals influences the extent to which individuals discriminate between congeners, which ultimately shapes mating outcomes (Andersson 1994). Therefore, it is important to understand how and why mating signals diverge between populations. Hybrid zones – regions where distinct species come into contact and interbreed – provide a natural experiment to examine the consequences of vocal divergence for behavioral isolation (Hewitt 1988).

Divergent mating signals can serve to reproductively isolate species with otherwise incomplete barriers to gene flow (Grant and Grant 1997; Price 2008). Learned vocalizations, such as oscine songs (Nottebohm 1972), have the potential to diverge rapidly via cultural evolution (Mason et al. 2016) and are therefore important pre-mating barriers to gene flow between hybridizing populations (Slabbekoorn and Smith 2002; Uy et al. 2018). In contrast, innate vocalizations diverge more slowly than learned vocalizations, and there is mixed evidence for the role of innate vocalizations as behavioral barriers to gene flow. For example, innate vocalizations serve as behavioral barriers to hybridization in *Alectoris* partridges (Ceugniet and Aubin 2001) and *Streptopelia* doves (De Kort et al. 2002), but not in *Callipepla* (Gee 2005), nor *Coturnix* quails (Derégnaucourt and Guyomarc’h 2003). This leaves an important gap in our understanding about innately derived vocalizations and how they vary between closely related species that are hybridizing.

In many bird species, both males and females vocalize (Odom and Benedict 2018). Vocal traits such as note length, complexity or production rate may differ between the sexes depending on their function in courtship and other behavioral contexts (Appleby et al. 1999; Odom and Mennill 2010; ten Cate 1997). For example, the sex that competes more for mates tends to vocalize more often (Sordahl 1979; Sung et al. 2005). Whereas male competition for mates is common across animals, females of some species are sex-role reversed, meaning they face stronger competition for mates than males do (Emlen and Oring 1977). Currently, we know very little about how female and male vocalizations compare in sex-role reversed species.

Sexually dimorphic vocalizations may also occur due to physical differences between males and females. In many species the sexes diverge in body size. As a consequence, morphological constraints on sound production can lead to distinctive spectral and temporal characteristics between male and female vocalizations (Ryan and Brenowitz 1985; Ten Cate 1997). Consistent with signal design theory, larger body size is often associated with lower sound frequencies both between and within sexes (Barbraud et al. 2000; Maurer et al. 2008). Some taxa are exceptions to this rule; for example, many female owls are larger than males but have higher frequency calls (Odom and Mennill 2010). Therefore, it is not clear whether signal design theory will hold in sex-role reversed species, in which females are often larger than males.

Jacanas are tropical, sex-role reversed shorebirds in which selection on females to compete for mates is stronger than on males (Jenni 1974). The Northern Jacana (*Jacana spinosa*) and Wattled Jacana (*J. jacana*) have been isolated for 700,000 years (Miller et al. 2014) and hybridize in a narrow region in Panama (Lipshutz et al. 2019). It is unknown whether their vocalizations are divergent, and what role their calls play in maintaining reproductive isolation between the species. Here, we quantify variation in temporal and spectral characteristics between the species and the sexes. We predict that vocalizations between *J. spinosa* and *J. jacana* will be divergent, and that the larger-bodied *J. spinosa* will have lower frequency-related characteristics. Second, we examine vocal divergence between males and females of both species. Jacanas have extreme sexual dimorphism in size, and females weigh up to 60% more than males (Emlen and Wrege 2004; Jenni and Collier 1972). Because female jacanas are larger than males, we predict that female vocalizations will have lower frequency-related characteristics. We also predicted that females should produce more calls than males, given that these species are sex-role reversed.

## Methods

### Sound recordings

We recorded vocalizations from June-August 2015 and June-July 2018 at 9 different sites in Panama (Fig. 1). Across these sites we recorded a total of 12 individuals of each species and sex. Birds were either stimulated with playback and a taxidermic mount to elicit vocalizations, or in some cases vocalizations were stimulated by the presence of the recordist near the bird’s territory.

**Figure 1.**
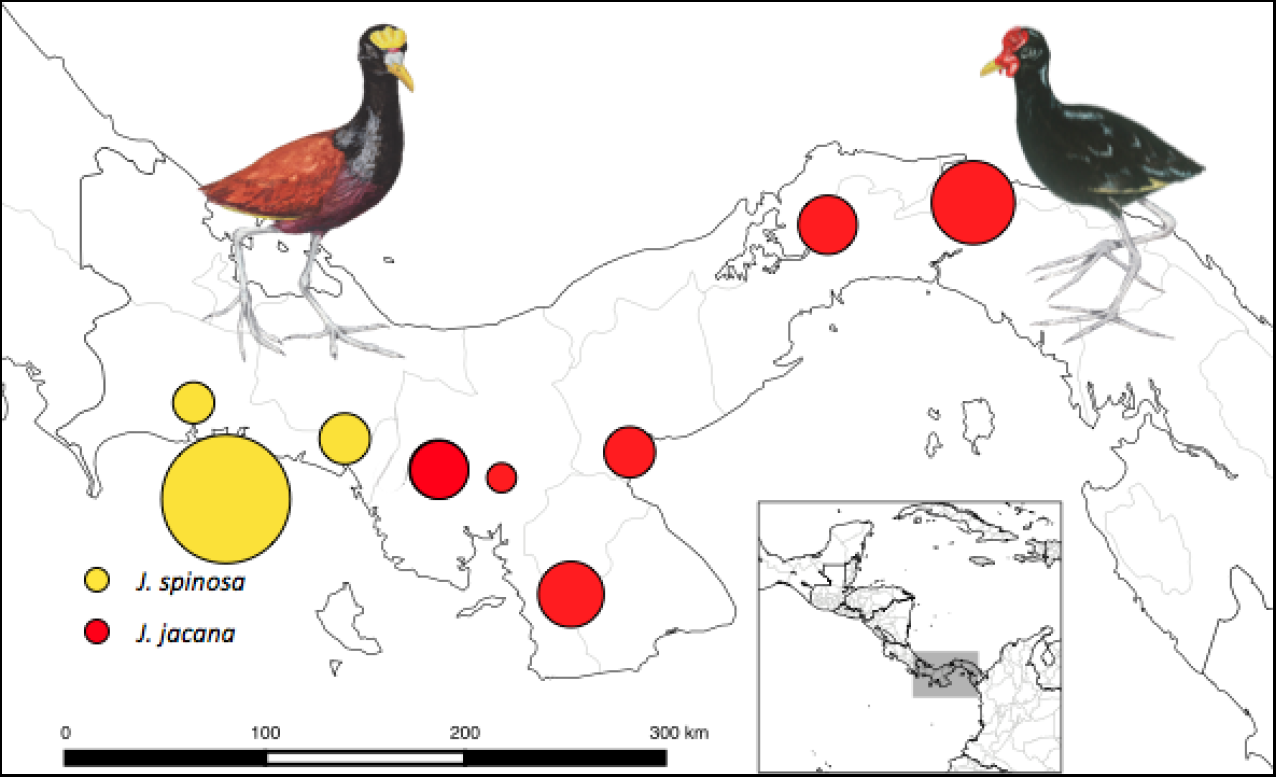
Sampling map of *Jacana spinosa* (yellow) and *J. jacana* (red) vocalizations recorded across the hybrid zone Panama. Circle size represents sample size (minimum 1, maximum 19).

Recordings were made using a Marantz PMD661 MKII solid state digital recorder (Marantz professional, Cumberland, Rhode Island, United States) set at 44.1 kHz sampling rate, 16-bit, and WAV file type, and a Sennheiser K6 power module with a Sennheiser M67 shotgun microphone and windscreen (Sennheiser electronic corporation, Wedemark, Germany). We divided continuous recordings for each individual into discrete call bouts (average = 6.1, range = call bouts per individual) in Audacity 2.1.2 (Audacity 2018). Call bouts are defined as a series of evenly spaced notes less than 1 second apart.

### Acoustic measurement

Jacana vocalizations contain harmonics covering a wide frequency bandwidth (Mace 1981). We took measurements on one call type, repeated note calls (Jenni 1974), as these were consistently found in recordings of both species and sexes (Fig. 2). We used the sound-analysis software Luscinia (Lachlan 2007) to generate Fourier-based spectrograms. Calls were high pass filtered to eliminate low frequency background noise below 200 Hz. We used the following settings to measure call variation: FF jump suppression = 20, Max. Frequency (Hz) = 15,000, Frame length (ms) = 5, Time step (ms) = 1, Spectrograph points = 221, Spectrogram Overlap % = 80, Dynamic range (dB) = 50, Dynamic equalization (ms) = 0, Dynamic comp. % = 100, Dereverberation % = 200, Dereverberation range (ms) = 100, Windowing function = Gaussian, Frequency zoom % = 150, Time zoom % = varies, Noise removal (dB) = 0, NR range1 (ms) = 50, R range2 (ms) = 50.

**Figure 2.**
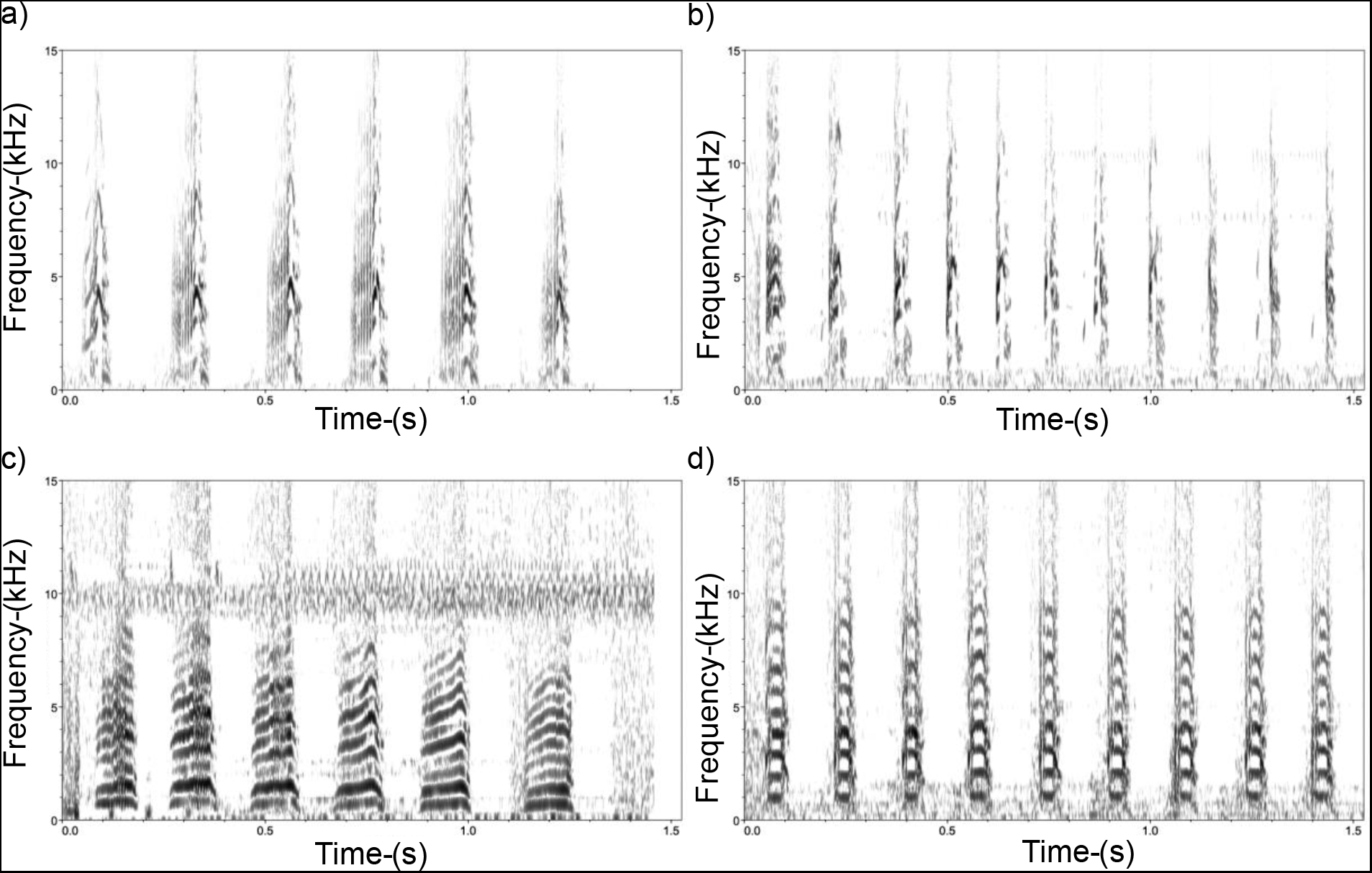
Spectrograms of vocalizations for a) female *J. spinosa*, b) male *J. spinosa*, c) female *J. jacana,* d) male *J. jacana*

We manually measured vocalizations in Luscinia (Lachlan 2007) by individually tracing each note. We used the Luscinia software to automatically calculate three acoustic parameters: note length (msec), peak frequency (frequency of the maximum amplitude, Hz), and fundamental frequency (Hz). We averaged these parameters for each note within a call bout (range 2-70 notes, mean 11.6) (Table 1). We also averaged the number of notes per bout for each individual to calculate a fourth acoustic parameter, notes per bout. The four parameters were averaged in Microsoft Excel after exporting from Luscinia.

**Table 1.** Sampling information and spectral and temporal characteristics (mean ± SE) for vocalizations recorded male and female *Jacana spinosa* and *J. jacana*.

|  | *Jacana spinosa* |  | *Jacana jacana* |  |
| --- | --- | --- | --- | --- |
|  | Male | Female | Male | Female |
| Number of Individuals | 12 | 12 | 12 | 12 |
| Notes per Bout | 15.97 ± 2.36 | 9.75 ± 1.55 | 14.03 ± 2.42 | 10.32 ± 1.66 |
| Range of Notes per Bout | 2 – 45 | 2 – 33 | 3 – 70 | 2 – 68 |
| Note Length (ms) | 70.17 ± 5.09 | 85.51 ± 7.52 | 78.12 ± 2.98 | 95.87 ± 4.3 |
| Peak Frequency (Hz) | 2898.36 ± 129.97 | 2539.14 ± 129.2 | 2221.52 ± 166.34 | 1521.63 ± 161.45 |
| Fundamental Frequency (Hz) | 1346.54 ± 112.46 | 1099.14 ± 99.4 | 731.83 ± 24.71 | 663.84 ± 23.46 |

### Statistical analysis

We compared the four call parameters between the species and sexes using Student’s *t*-tests or Wilcoxon rank sum, depending on whether the parameters were normally distributed or not. We also summarized the four acoustic parameters with a principal components analysis (PCA) using the prcomp function in R version 3.3.2 (R-Core-Team, 2015). Prior to the PCA, we log-transformed acoustic data to fulfill assumptions of multi-normality. We retained two PC scores that explained 60.6% and 19.7% of the variation in the acoustic parameters, totaling 80.7% cumulative proportion of variation (Table S1). Frequency variables loaded negatively onto PC1 and positively onto PC2. Note length loaded positively on PC1 and PC2, and notes per bout loaded negatively onto PC1 and PC2. We compared vocalizations between the species and sexes using linear mixed effects models with the nmle package (Pinheiro et al. 2018) in R. We included vocalization PC1 and PC2 as separate response variables, species and sex as the fixed effects, and site as a random effect. We visually inspected residual plots to ensure they did not deviate from normality. We used a type III ANOVA to determine whether species and/or sex were significant predictors of variation in the PCs.

We also conducted a discriminant function analysis (DFA) in R using the package flipMultivariates to assess whether spectral or temporal parameters could distinguish between species and sexes (https://github.com/Displayr/flipMultivariates/).

## Results

### Species differences in vocalizations

Vocalizations are different between these two jacana species, particularly regarding spectral characteristics (Table 1; Fig. 3). For both males and females, *J. spinosa* calls have significantly higher peak (t = −5.1, df = 41.8, *P* < 0.001) and fundamental frequencies (W = 42, *P* < 0.001) than *J. jacana* calls. Species was also a significant predictor both of vocalization PC1 (*F*_1,7_ = 24.8, *P* = 0.002) and PC2 (*F*_1,7_ = 16.7, *P* = 0.005) (Fig. S1). Using all acoustic parameters, 44 out of 48 (92%) individuals were classified to the correct species by a DFA. The best variables to distinguish the species were peak frequency (*r*^2^ = 0.36, *P* < 0.001) and fundamental frequency (*r*^2^= 0.48, *P* < 0.001).

**Figure 3.**
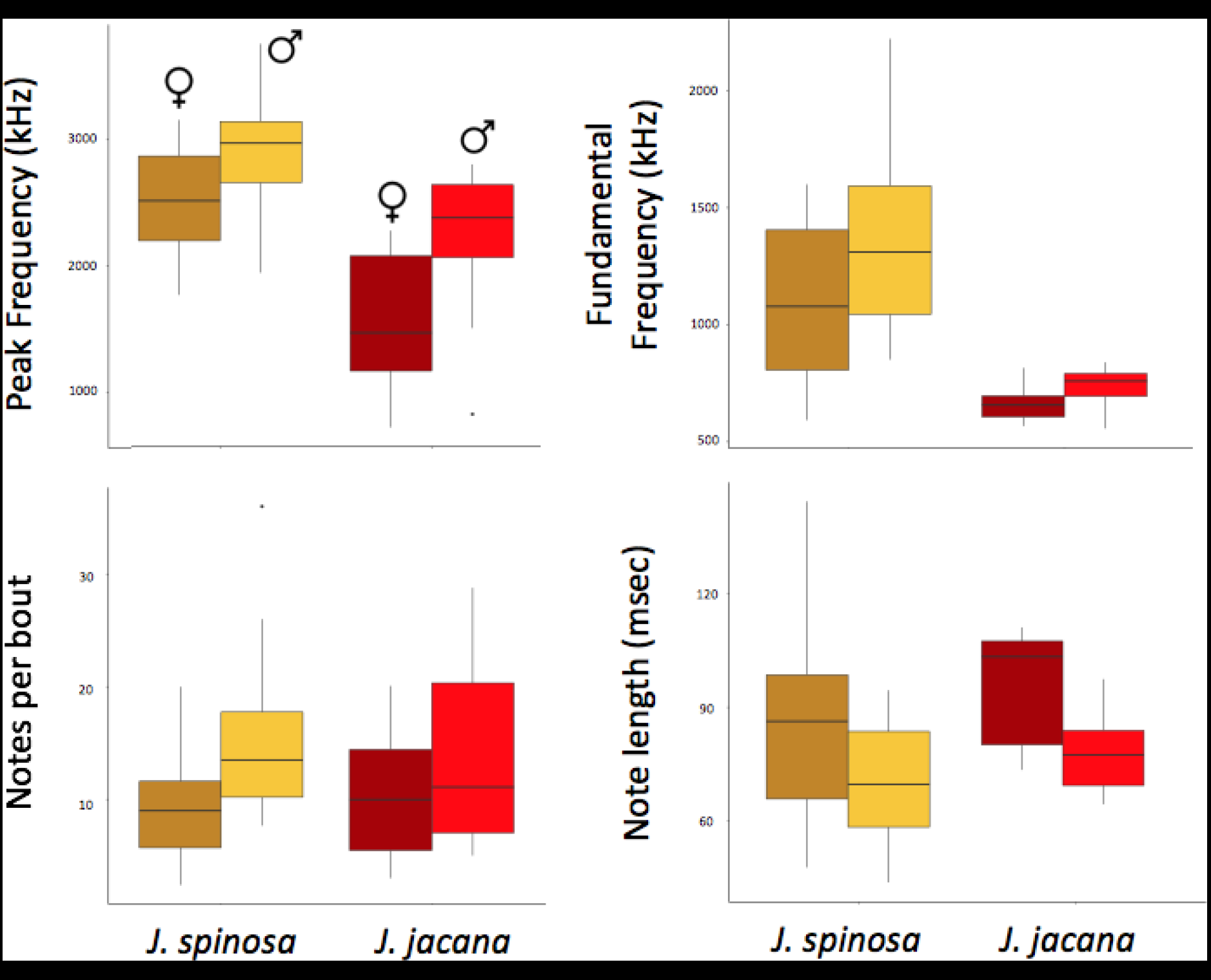
Mean spectral and temporal characteristics for vocalizations recorded from male (light) and female (dark) *Jacana spinosa* (yellow) and *J. jacana* (red).

### Sex differences in vocalizations

Vocalizations are also different between males and females of both species (Table 1; Fig. 3). When species are combined, males have significantly more calls within a bout (W = 189.5, *P* = 0.043) and higher peak frequency calls (t = −2.7, df = 44.9, *P* = 0.009) than females. Females have longer calls than males (t = 3.1, df = 40.7, *P* = 0.003). Sex was also a significant predictor of vocalization PC1 (*F*_1,38_ = 14.8, *P* = < 0.001) but not of PC2 (Fig. S1). Using all acoustic parameters, 45 out of 48 (73%) individuals were classified to the correct sex by a DFA. The best variables to distinguish the sexes were call length (*r*^2^ = 0.17, *P* = 0.005), peak frequency (*r*^2^ = 0.14, *P* = 0.014), and calls within a bout (*r*^2^ = 0.12, *P* = 0.028).

The sexes were more strongly differentiated by peak frequency in *J. jacana* (t = −3.0193, df = 21.98, *P* = 0.006) than in *J. spinosa* (t = −1.9602, df = 21.999, *P* = 0.063). Similarly, males and females differed more strongly in note length in *J. jacana* (t = 3.3939, df = 19.602, *P* = 0.003) than in *J. spinosa* (t = 1.6899, df = 19.318, *P* = 0.11). In contrast, the sexes differed more strongly in calls within a bout for *J. spinosa* (W = 38.5, *P* = 0.057) than *J. jacana* (W = 53, *P* = 0.29).

## Discussion

*Jacana spinosa* and *J. jacana* calls are different - *J. spinosa* calls are significantly higher in peak and fundamental frequency. The sexes are consistently different across both species, such that male calls have higher peak frequency and more notes than female calls, whereas female calls have longer note lengths.

### Species Differences

Vocalizations of the two species of jacana have diverged spectrally but are similar temporally. In other species of non-oscine birds that do not learn their songs, temporal traits change at slower rates than frequency-related traits (Miller and Baker 2009, Seneviratne et al. 2012), which is consistent with our findings that spectral traits were more divergent than temporal traits between these jacana species. Counter to the prediction that smaller-bodied species should have calls with higher frequencies (Ryan and Brenowitz 1985), larger-bodied *J. spinosa* have higher peak and fundamental frequency vocalizations than smaller-bodied *J. jacana*, for both sexes. One hypothesis for this contradiction is that species divergence in vocalizations could relate to differing environmental or habitat characteristics that have shaped their call frequencies (Morton 1975, Endler 1992). In a prior study, species distribution modeling indicated that *J. spinosa* favors a warmer and wetter environment (Miller et al. 2014), suggesting that the species have diverged in habitat preferences. Future work could compare the vegetative cover in the habitat of each species to determine whether the higher frequency vocalizations of *J. spinosa* relate to a more open habitat. Another potential explanation for vocal divergence could be a difference in the syringeal or bill structure between species (Seneviratne et al. 2012; Kingsley et al. 2018).

### Sex Differences

We found that frequency differences in jacana vocalizations matched our predictions for females and males based on their body size dimorphism; males of both species produce higher frequency calls than the larger-bodied females. In many species with sexually dimorphic body size, the smaller of the two sexes produces higher frequency vocalizations (Maurer et al. 2008). Female-biased size dimorphism is common in shorebirds other than jacanas, and larger females also have lower frequency calls (Heidemann and Oring 1976; Douglas 1998). Contrary to our expectation, male jacanas within our study also emitted more notes per calling bout than females. One likely explanation for this is mate attraction: in the polyandrous Bronze-winged Jacana (*Metopidius indicus*), individual males that called more frequently received more copulations than other co-mates (Butchart et al. 1999). Calling rate could be a sexually selected trait in male Neotropical jacanas, and this hypothesis should be tested using behavioral playback experiments.

### Application and conclusion

We found that Neotropical jacanas diverged significantly in the peak and fundamental frequencies of their vocalizations. Diverged vocal signals could promote reproductive isolation between the two species when they come into contact. Furthermore, these spectral characteristics differed between the species for both sexes, suggesting that both male and female signals could facilitate species-specific discrimination in the hybrid zone. A phenotypic and genomic analysis of the jacana hybrid zone found that species-specific traits such as plumage and facial ornamentation were likely prezygotic barriers that maintain species boundaries (Lipshutz et al. 2019). Future playback studies could assess the relative role of visual and vocal signals as behavioral barriers to mating between the species. This phenotypic differentiation between *J. spinosa* and *J. jacana* likely contributes to the low occurrence of hybrids within the narrow hybrid zone and may be one of the reasons for limited hybridization between the species.

## Acknowledgments

This material is supported by the University of Tennessee’s Ready for the World Chancellor’s Honors Program and Summer Undergraduate Research Internship to EJB, National Science Foundation Graduate Research Fellowship Grant No. 1154145, Doctoral Dissertation Improvement Grant IOS‐1818235, and a Smithsonian Tropical Research Institute short-term fellowship to SEL, Tulane University’s CELT summer research and faculty-student scholarly engagement grants to TB, Tulane University’s Newcomb Scholars and Dean’s summer research grants to GZ, and a Louisiana Board of Regents NSF EPSCoR LINK Grant No. 177 to EPD. Any opinion, findings, and conclusions or recommendations expressed in this material are those of the authors(s) and do not necessarily reflect the views of the National Science Foundation. Scientific recording in Panama is done with the prior approval of MiAmbiente (formerly ANAM), Panama's environmental authority (permit numbers: SE/A-45-12, SE/A-46-14, SE/A-17-18), and recording techniques were approved by the Institutional Animal Care and Use Committee of the Smithsonian Tropical Research Institute (IACUC permits: 2012-0315-2015, 2018-0116-2021), Tulane University (IACUC permit: 0446R), and the University of Tennessee (IACUC permit: 2573). We thank the STRI Bird Collection for preparing taxidermic mounts. Daniel Ernesto Buitrago and Marcelo Araya Salas assisted with translating the abstract into Spanish. Jacana illustrations by Stephanie McClelland.

## Data accessibility

Recordings are available on xeno canto. *Jacana spinosa*: https://www.xeno-canto.org/set/4909

---

**Supplemental Figure 1.**
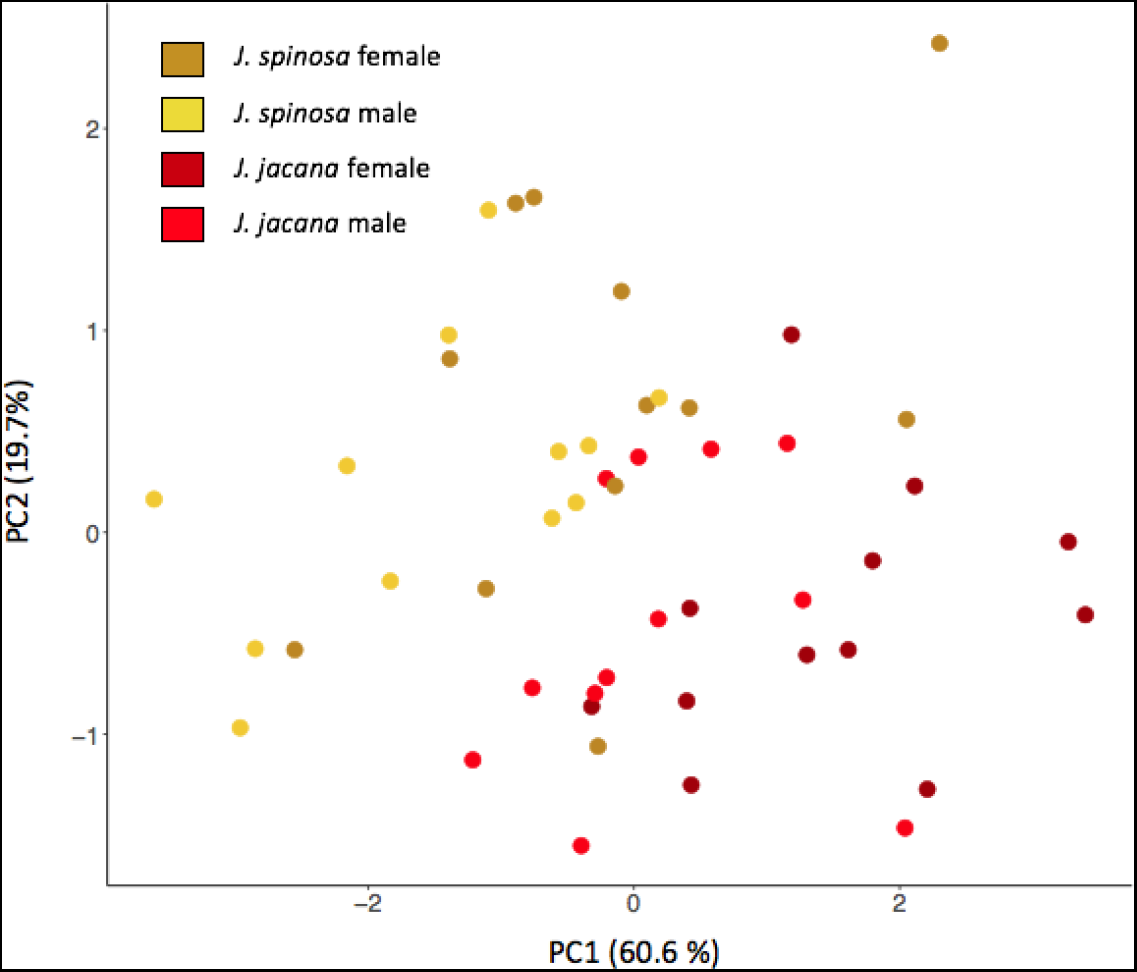
Principal components analysis of *Jacana spinosa* (yellow) and *Jacana jacana* (red) males (light) and female (dark) vocalizations.

**Supplemental Table 1.**
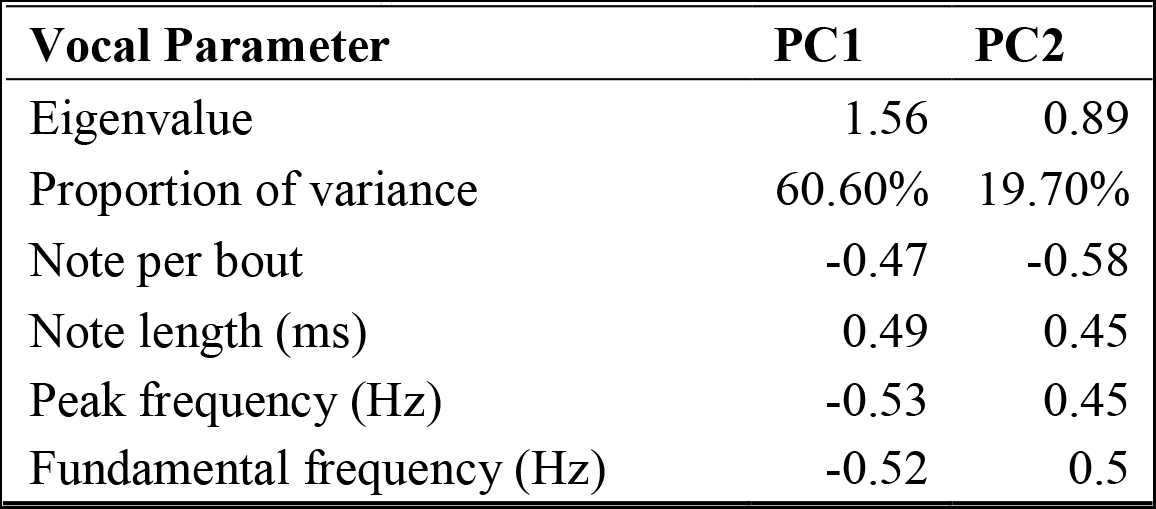
Loadings for Principal Component Analysis

## Literature Cited

Andersson M. 1994. Sexual selection (Monographs in behavior and ecology). Princeton (NJ): Princeton University Press.

Appleby BM, Yamaguchi N, Johnson PJ, MacDonald DW. 1999. Sex-specific territorial responses in Tawny Owls Strix aluco. Ibis. 141:91–99.

Audacity. 2018. Audacity ^®^ | Free, open source, cross-platform audio software for multi-track recording and editing. Audacity. doi: ISSN 1980-4431.

Baker MC, Boylan JT. 1999. Singing Behavior, Mating Associations and Reproductive Success in a Population of Hybridizing Lazuli and Indigo Buntings. The Condor. 101:493–504.

Barbraud C, Mariani A, Jouventin P. 2000. Variation in call properties of the snow petrel, Pagodroma nivea, in relation to sex and body size. Australian Journal of Zoology. 48:421–430.

Butchart S, Seddon N, Ekstrom JMM. 1999. Yelling for sex: Harem males compete for female access in bronze-winged jacanas. Animal Behaviour. 57:637–646.

Catchpole CK, Slater PJB. 2008. Bird song: Biological themes and variations, second edition. Cambridge (UK): Cambridge University Press.

Ceugniet M, Aubin T. 2001. The rally call recognition in males of two hybridizing partridge species, red-legged (Alectoris rufa) and rock (A. graeca) partridges. Behavioural Processes. 55:1–12.

Coyne JA, Orr AH. 2004. Speciation. Sunderland (MA): Sinauer Associates.

De Kort SR, Den Hartog PM, Ten Cate C. 2002. Vocal signals, isolation and hybridization in the vinaceous dove (Streptopelia vinacea) and the ring-necked dove (S. capicola). Behavioral Ecology and Sociobiology. 51:378–385.

Derégnaucourt S, Guyomarch JC. 2003. Mating call discrimination in female European (Coturnix c. coturnix) and Japanese quail (Coturnix c. japonica). Ethology. 109:107–119.

Douglass HD. 1998. Response of Eastern Willets (Catoptrophorus s. semipalmatus) to Vocalizations of Eastern and Western (C. s. inornatus) Willets. The Auk. 115:514–518.

Emlen ST, Oring LW. 1977. Ecology, sexual selection, and the evolution of mating systems. Science. 197:215–223.

Emlen ST, Wrege PH. 2004. Size Dimorphism, Intrasexual Competition, and Sexual Section in Wattled Jacana (Jacana jacana), a Sex-Role-Reversed Shorebird in Panama. The Auk. 121:391–403.

Endler J. 1992. Signals, Signal Conditions, and the Direction of Evolution. The American Naturalist 139:S125.

Gee JM. 2005. No species barrier by call in an avian hybrid zone between California and Gambels quail (Callipepla californica and C. gambelii). Biological Journal of the Linnean Society. 86:253–264.

Grant PR, Grant BR. 1997. Genetics and the origin of bird species. Proceedings of the National Academy of Sciences. 94:7768–7775.

Hunt J, Breuker CJ, Sadowski JA, Moore AJ. 2009. Male-male competition, female mate choice and their interaction: Determining total sexual selection. Journal of Evolutionary Biology. 22:13–26.

Heidemann MK, Oring LW. 1976. Functional Analysis of Spotted Sandpiper (Actitis macularia) Song. Behaviour. 56:181–193.

Irwin DE, Price T. 1999. Sexual imprinting, learning and speciation, Heredity. 82:347–354.

Jenni DA. 1974. Evolution of polyandry in birds. Integrative and Comparative Biology. 14:129–144.

Jenni DA, Collier G. 1972. Polyandry in the American Jaçana (Jacana spinosa). The Auk. 89:743–765.

Kingsley EP, Eliason CM, Riede T, Li Z, Hiscock TW, Farnsworth M, Thomson SL, Goller F, Tabin CJ, Clarke JA. 2018. Identity and novelty in the avian syrinx. Proceedings of the National Academy of Sciences. 115:10209–10217.

Konishi M. 1963. The Role of Auditory Feedback in the Vocal Behavior of the Domestic Fowl. Zeitschrift für Tierpsychologie. 20:349–367.

Lachlan RF. 2007. Luscinia: a bioacoustics analysis computer program.

Lipshutz SE, Meier JI, Derryberry GE, Miller MJ, Seehausen O, Derryberry EP. 2019. Differential introgression of a female competitive trait in a hybrid zone between sex-role reversed species. Evolution. 73:188–201.

Lynch A. 1996. The population memetics of birdsong. In: Kroodsma DE, Miller EH, editors. Ecology and evolution of acoustic communication in birds. Ithaca (NY): Comstock Publishers; p.181–197.

Mace TR. 1981. Causation, function, and variation of the vocalizations of the Northern Jacana, Jacana spinosa. [dissertation]. Missoula (MT): University of Montana.

Mason NA, Burns KJ, Tobias JA, Claramunt S, Seddon N, Derryberry EP. 2016. Song evolution, speciation, and vocal learning in passerine birds. Evolution. 71:786–796.

Maurer G, Smith C, Süsser M, Magrath RD. 2008. Solo and duet calling in the pheasant coucal: Sex and individual call differences in a nesting cuckoo with reversed size dimorphism. Australian Journal of Zoology. 56:143–149.

Miller EH, Baker AJ. 2009. Antiquity of Shorebird Acoustic Displays. The Auk. 126:454–459.

Miller MJ, Lipshutz SE, Smith NG, Bermingham E. 2014. Genetic and phenotypic characterization of a hybrid zone between polyandrous Northern and Wattled Jacanas in Western Panama. BMC Evolutionary Biology. 14:227.

Nottebohm F. 1972. The Origins of Vocal Learning. The American Naturalist. 106:116–140.

Odom, KJ, Benedict L. 2018. A call to document female bird songs: Applications for diverse fields. The Auk 135:314–325.

Odom KJ, Mennill DJ. 2010. A Quantitative Description of the Vocalizations and Vocal Activity of the Barred Owl. The Condor. 112:549–560.

Pinheiro J, Bates D, DebRoy S, Sarkar D, Heisterkamp S, Van Willigen B. 2018. nlme: Linear and Nonlinear Mixed Effects Models. R package version 3.1-137, Retrieved from https://cran.r-project.org/package=nlme.

Price T. 2008. Speciation in Birds. Greenwood Village (CO): Roberts and Company Publishers.

R-Core-Team. 2015. R: A language and environment for statistical computing. Vienna, Austria: R Foundation for Statistical Computing.

Ryan MJ, Brenowitz EA. 1985. The Role of Body Size, Phylogeny, and Ambient Noise in the Evolution of Bird Song. The American Naturalist. 126:87–100.

Seneviratne SS, Jones IL, Carr SM. 2012. Patterns of vocal divergence in a group of non-oscine birds (auklets; Alcidae, Charadriiformes), Evolutionary Ecology Research. 1:95–112.

Slabbekoorn H, Smith TB. 2002. Bird song, ecology and speciation, Philosophical Transactions of the Royal Society B: Biological Sciences. 357:493–503.

Sordahl TA. 1979. Vocalizations and behavior of the Willet Catoptrophorus semipalmatus. Wilson Bulletin. 91:551–574.

Sung H-C, Miller EH, Flemming SP. 2005. Breeding vocalizations of the piping plover (Charadrius melodus): structure, diversity, and repertoire organization. Canadian Journal of Zoology. 83.579–595.

Taoka M, Sato T, Kamada T, Okumura H, 1989. Sexual Dimorphism of Chatter-Calls and Vocal Sex Recognition in Leachs Storm-Petrels (Oceanodroma leucorhoa). The Auk: Ornithological Advances. 106:498–501.

Ten Cate C. 1997. Sex Differences in the Vocalizations and Syrinx of the Collared Dove (Streptopelia decaocto). The Auk. 114:22–39.

Uy JAC, Irwin DE, Webster MS. 2018. Behavioral Isolation and Incipient Speciation in Birds. Annual Review of Ecology, Evolution, and Systematics. 49:1–24.

